# Integrative analysis of the hydroxypyruvate reductases revealing their distinct roles in photorespiration of Chlamydomonas

**DOI:** 10.1101/2021.04.20.440714

**Authors:** Menglin Shi, Lei Zhao, Yong Wang

## Abstract

Photorespiration plays an important role in maintaining normal physiological metabolism in higher plants and other oxygenic organisms such as algae. The unicellular eukaryotic organism *Chlamydomonas* is reported to have a different photorespiration system from that in higher plants, and only two out of nine genes encoding photorespiratory enzymes have been experimentally characterized. Hydroxypyruvate reductase (HPR), which is responsible for the conversion of hydroxypyruvate into glycerate, is poorly understood and not yet explored in Chlamydomonas. To identify the candidate genes encoding hydroxypyruvate reductase in Chlamydomonas (CrHPR) and uncover their elusive functions, we performed sequence comparison, enzyme activity measurement, subcellular localization, and analysis of knockout/knockdown strains. Together we identify five proteins to be good candidates as CrHPRs, all of which are detected with the activity of hydroxypyruvate reductase. CrHPR1, a NADH-dependent enzyme in mitochondria, may function as the major component of photorespiration, and deletion of *CrHPR1* causes severe photorespiratory defects. CrHPR2 takes parts in the cytosolic bypass of photorespiration as the compensatory pathway of CrHPR1 for the reduction of hydroxypyruvate. CrHPR4, with NADH as the cofactor, may participate in photorespiration by acting as the chloroplastidial glyoxylate reductase in glycolate-quinone oxidoreductase system. Therefore, our results reveal that the CrHPRs are far more complex than previously recognized, and provide a greatly expanded knowledge base for studies to understand how CrHPRs perform their functions in photorespiration. These will facilitate the genetic engineering for crop improvement by synthetic biology.

**Brief summary:** Identification and characterization of genes encoding hydroxypyruvate reductases in Chlamydomonas, demonstrating difference in the enzymatic activity, subcellular location, as well as function in photorespiration.

## INTRODUCTION

Photosynthesis and photorespiration are the two major pathways of plant primary metabolism mediated by the bifunctional enzyme ribulose bisphosphate carboxylase/oxygenase (Rubisco, Griffiths, 2006). Since the cellular O_2_ concentration is much higher than that of CO_2_, correspondingly high amount of photosynthetically fixed carbon is released via the oxidation of Rubisco during photorespiration (Somerville, 2001). The loss of fixed carbon associated with photorespiration could be greatly elevated by the rising temperature and drought, resulting in the severe reduction of crop yields (Bauwe et al., 2012). Hence, photorespiration is becoming the important target for crop improvement and obtaining more attention considering that the world will face more serious challenges such as extreme climate and severe food shortage (Ort et al., 2015).

Photorespiration, conserved in higher plants and most algae, comprises a series of nine consecutive enzymatic reactions distributed over chloroplasts, peroxisomes, mitochondria, and the cytosol (Bauwe et al., 2012). It starts from the catalysis mediated by Rubisco in chloroplast, in which ribulose 1,5-bisphosphate (RuBP) is oxidized into 2-phosphoglycolate (2-PG) and 3-phosphoglycerate (3-PGA, Tcherkez, 2013). 2-PG is dephosphorylated by the phosphoglycolate phosphatase, leading to the generation of glycolate which enters photorespiratory metabolism flux for the further conversion (Douce and Heldt, 2000). In the penultimate step of photorespiration cycle, hydroxypyruvate is converted to glycerate catalyzed by hydroxypyruvate reductase (HPR), and eventually to Calvin-Benson intermediate 3-PGA (Hagemann and Bauwe, 2016). Despite the good consensus on the photorespiratory cycle, the serial catalytic enzymes and the detailed functions till now have not yet been globally characterized (Eisenhut et al., 2019). One question that remains to be resolved is concerning the presence and function of hydroxypyruvate reductase in *Chlamydomonas* photorespiration, which is the topics of the present study.

HPRs are highly conserved evolutionarily, however, they are only experimentally characterized in limited extent, especially in terms of underlying gene at molecular level. Since 1970s, HPRs have been purified and studied on their enzymological property with then state-of-the-art methodology, and the studies mainly focused on HPRs from barley (Kleczkowski et al., 1990), Spinach (Kleczkowski et al., 1988, 1991), Cucumber (Titus et al., 1983; Greenler et al., 1989; Schwartz et al., 1992), Pumpkin (Hayashi et al., 1996; Mano er al., 2000), Arabidopsis (Mano et al., 1997; Cousins et al., 2008) and Chlamydomonas (Stabenau et al., 1974; Husic and Tolbert, 1987). Until recently, functional details of the underlying genes are revealed in model organism like *Arabidopsis*. The deletion of *HPR1* causes no visible alternation of growth or photorespiration in atmospheric air, differentiating from the lethal phenotype displayed by the mutants with impairment in other photorespiratory components (Murray et al., 1989; Timm et al., 2008; Cousins et al., 2011; Ye et al., 2014). The non-typical photorespiratory phenotypes of *hpr1* could be explained by the study from *Arabidopsis*, in which the missing of peroxisome*-*targeting AtHPR1 is partially compensated by the cytosolic bypass of photorespiration mediated by AtHPR2 (Timm et al., 2008; Li et al., 2019). Nevertheless, the combined deletion of both *HPR1* and *HPR2* does not result in the lethal phenotype of *hpr1hpr2*, though the typical photorespiratory characteristics were detected (Timm et al., 2008; Ye et al., 2014), suggesting the presence of additional HP-reducing enzyme. With BLAST search, HPR3 was identified to be the potential candidate for such enzyme in *Arabidopsis*, and it showed activity with HP using NADPH as the co-substrate (Timm et al., 2011). Interestingly, HPR3 could accept glyoxylate as a substrate, thus it may represent an additional bypass both to known HPRs and to glyoxylate reductases in chloroplasts (Timm et al., 2011).

Unlike the higher plants, *Chlamydomonas* employed a CO_2_ concentration mechanism (CCM) to increase CO_2_ concentration in the vicinity of Rubisco, and its photorespiration system is assumed to work differently (Wang et al., 2015). Shortly after transfer from high CO_2_ to low CO_2_, but before the induction of CCM, photorespiration metabolism was induced rapidly and briefly (Brueggeman et al., 2012). Moreover, photorespiration is assumed to pass through the mitochondria rather than peroxisome in *Chlamydomonas*, which is not yet to be globally studied and confirmed (Nakamura et al., 2005). In terms of HPR, Stabenau initially detected the CrHPR activity in mitochondria (Stabenau, 1974), then Husic and Tolbert further investigated the enzymatic characteristics with the cell extracts (1985). In the decades since then, no further research on CrHPRs is reported, and both the underlying genes and their detailed functions remain waiting to be discovered.

To search for the elusive CrHPRs, we have now exploited the Chlamydomonas genome and identified 5 proteins as candidates for being novel CrHPRs. Detailed studies indicate that all five CrHPRs are detected with the activity of HPR. Using both bioinformatics and cell biology approaches, we were able to define the subcellular location of CrHPRs. We further addressed the physiological roles of *CrHPR1, CrHPR2* and *CrHPR4* by analyzing the respective knockout/knockdown strains, and revealed their distinct roles in photorespiration. These findings are important steps toward understanding how CrHPRs perform their functions, and will facilitate the genetic engineering for crop improvement by providing prime targets for modification.

## RESULTS

### Identification and bioinformatic analysis of CrHPRs

HPR proteins are highly conserved over a wide range of species (Kutner et al., 2018). The putative Chlamydomonas HPR homolog is the protein encoded by *Cre06*.*g295450*.*t1*.*2* (hereafter designated as *CrHPR1*). Sequence alignment of HPR proteins from select species indicated the conservation of CrHPR1 as hydroxypyruvate reductase, manifesting as the presence of highly conserved NAD(P)-binding motif, NAD recognition sites, substrate-orienting and catalytic pair domains (Fig. S1).

To identify all candidate CrHPRs, protein sequence of CrHPR1 was used to search against the Chlamydomonas Proteome via the BLASTP function built on the widely used online service, including Phytozome 12, HMMER or AlgaePath, with the default parameters. The same list of eight proteins was retrieved with each of the three independent searches, and CrHPR1 itself was identified with the lowest E-value. Three remarkably similar proteins encoded by *Cre07*.*g344400, Cre07*.*g344550* and *Cre07*.*g344600*, respectively and annotated as phosphoglycerate dehydrogenase, were identified possibly because of the presence of NAD-binding and catalytic domains. Since they were reported to function in thylakoid membrane remodeling in response to adverse environmental conditions (Du et al., 2018), they are unlikely candidates for CrHPRs related to photorespiration, and were not characterized further.

To confirm whether the proteins encoded by *Cre01*.*g019100*.*t1*.*2, Cre02*.*g087300*.*t1*.*3, Cre07*.*g324550*.*t1*.*3* and *Cre16*.*g689700*.*t1*.*3*, were authentic CrHPRs, sequence alignment was performed with CrHPR1 as the reference. As shown in Fig. S2, these five proteins shared the conserved HPR domains suggesting that they are good candidates as hydroxypyruvate reductase. Therefore, *Cre01*.*g019100*.*t1*.*2,Cre02*.*g087300*.*t1*.*3, Cre07*.*g324550*.*t1*.*3 and Cre16*.*g689700*.*t1*.*3* were serially named as *CrHPR2, CrHPR3, CrHPR4 and CrHPR5* according to their location in chromosome, respectively, which together with *CrHPR1* made of five candidate genes encoding hydroxypyruvate reductase in Chlamydomonas genome.

### CrHPRs are proteins with hydroxypyruvate reductase activity

To determine how reliable our identification of potentially novel CrHPRs was and compare their enzymatic properties, CrHPRs identified above were heterologous expressed in *E. coli* and tested different substrate and cofactor combinations (Table 1). The expression and purification of tagged-recombinant CrHPRs were assayed with SDS-PAGE gels. As shown in Fig. S3, the recombinant CrHPR proteins were obtained at high purity, and each migrated at the expected molecular weight on the gels.

**Table 1.** Activity assay of recombinant CrHPRs with substrate and cofactor combinations.

| Substrates | CrHPR1 | CrHPR2 | CrHPR3 | CrHPR4 | CrHPR5 |
| --- | --- | --- | --- | --- | --- |
| Hydroxypyruvate:NADH | 312.5±19.3 | 2.58±0.01 | - | 46.23±1.31 | 1.86±0.11 |
| Hydroxypyruvate:NADPH | 1.70±0.24 | 11.03±0.57 | 3.03±0.01 | - | 292.40±13.49 |
| Glyoxylate:NADH | 13.91±0.48 | 2.05±0.08 | - | 14.35±0.20 | 0.91±0.04 |
| Glyoxylate:NADPH | 0.14±0.02 | 3.66±0.09 | 5.65±0.01 | - | 5.72±0.19 |
| Pyruvate:NADH | - | - | - | 666.67±12.36 | - |
| Pyruvate:NADPH | - | - | - | - | - |

Shown are mean activities ± SD from three independent measurements with tag-purified recombinant CrHPRs ( $\mu\text{mol}\cdot\text{min}^{-1}\cdot\text{mg}^{-1}$ protein).

With the purified recombinant CrHPRs, we measured the enzymatic activity with different substrate and cofactor combinations (Table 1). Tag-purified recombinant CrHRP1 was active in the presence of NADH and hydroxypyruvate, whereas NADPH-dependent rate was >180 times lower (312.5 *vs* 1.70 μmol·min^-1^ ·mg ^-1^ protein). The enzyme also accepted glyoxylate as a substrate, but at a considerably lower efficiency in the presence of NADH (312.5 *vs* 13.91 μmol·min^-1^ ·mg ^-1^ protein). These parameters established CrHPR1 as the NADH-dependent HPR in *Chlamydomonas*. By contrast, CrHPR5 mainly presented as the NADPH-dependent HPR with a much higher enzymatic activity than that of any other combinations (>50 times).

CrHPR3 and CrHPR4 were detected with cofactors specificity, and they were only active in the presence of NADPH and NADH, respectively. Intriguingly, CrHPR4 could accept pyruvate as substrate with correspondingly high enzymatic activity, suggesting its potential roles in fermentative metabolism (Burgess et al., 2015). Although it also accepted both hydroxypyruvate and glyoxylate as substrates, the catalytic efficiency is much lower than that with pyruvate (<15 times). The purified CrHPR2 showed relaxed activity with both NADH and NADPH, and hydroxypyruvate was the more favored substrate than glyoxylate.

### CrHPRs may function in multiple subcellular regions

To explore the potential functions of CrHPRs, phylogenetic tree was generated to infer their evolution characteristics using the maximum likelihood algorithm (Kumar et al., 2015). As shown in Fig. S4, CrHPR1 was assigned into the plant subgroup, suggesting its conservation in plant-specific metabolism during evolution, and it’s consistent with the alignment analysis (Fig. S1). CrHPR4 presented closely association with the photosynthetic cyanobacteria subgroup, which may imply its potential role in chloroplast considering the endosymbiotic hypothesis (Martin et al., 2015). CrHPR2, CrHPR3 and CrHPR5 are likely originated from the bacteria as they were put into the ascomycetes and firmicutes subgroups, suggesting that they may participate in mitochondrial or cytosolic metabolism.

To determine the reliability of subcellular localization inferred from phylogenetic analysis, either the N-terminal or C-terminal of CrHPRs was tagged with mCerulean fluorescent protein (CFP) as described in methods. Then the constructs were transformed into WT/CC-125 strain, screened for the positive clones, and assayed by laser confocal microscopy. As visible in Fig.S5, CrHPR1- and CrHPR5-CFP (amino acids of the N-terminus of CrHPR2 and CrHPR5) accumulated in the punctuated dots distributed in cytosol, and they likely function in mitochondria as no putative peroxisomal localization sequence, PTS1 or PTS2, were identified in their sequences (Gould et al., 1987; Swinkels et al., 1991). Using both N-terminal and C-terminal tagging only, CrHPR2 and CrHPR3 were confirmed to diffusely presented in the cytosol.

Chloroplast targeting was found for CrHPR4 (amino acids of the C-terminus) supported by the overlapping between CFP-CrHPR4 signal and chlorophyll autofluorescence, and it’s in accordance with previous study (Burgess et al., 2015). Therefore, inference of CrHPRs’ subcellular localization from phylogenetic analysis is in consistent with the results of fluorescence microscopy.

### NADH-dependent CrHPRs take parts in photorespiration as the major components

Previous studies revealed that, cofactors, such as NADH or NADPH, are combined with the substrate in the catalytic reaction of HPRs (Lassalle et al., 2016). Based on this, we designed a strategy of two steps to determine the CrHPRs participating in photorespiration.

Step 1: We examined the cofactors (NADH or NADPH) employed in photorespiration, from which we could infer the related CrHPRs on the basis of their catalytic characteristics (Table 1). As shown in Fig. 1A, the enzyme activity of NADH-dependent HPRs was much higher in photorespiration induced by air condition than that under non-photorespiration condition (CO_2_ condition) while the activity of NADPH-dependent HPRs was only slightly changed, which demonstrated the major roles of NADH-dependent CrHPRs in photorespiration. Since CrHPR1, CrHPR2, CrHPR4 and CrHPR5 were detected with the activity of HPRs in the presence of NADH (Table 1), thus their participation in photorespiration were examined further.

**Fig. 1.**
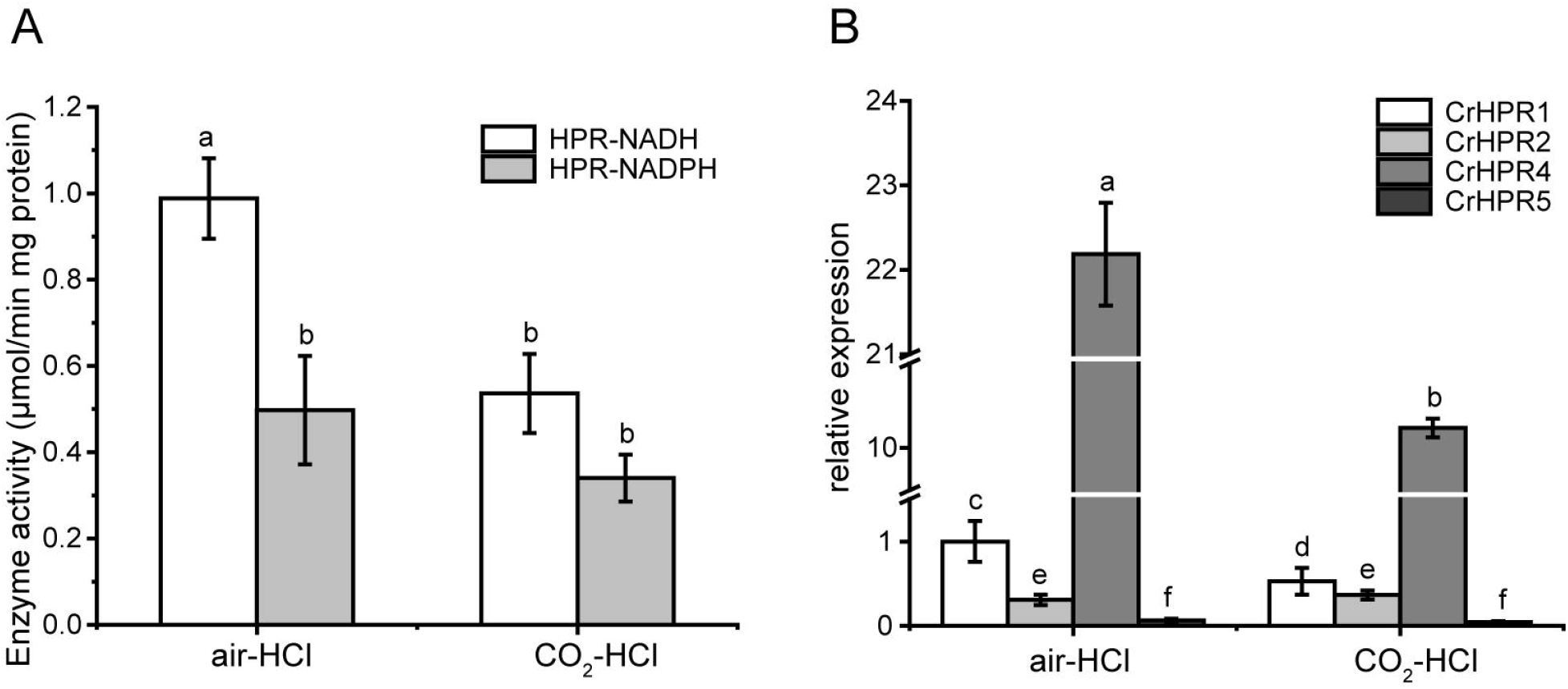
Enzyme activity assay and measurement of CrHPRs transcripts in photorespiration (air) and non-photorespiration conditions (CO_2_) (A) Assay of CrHPRs enzyme activity. (B) Detection of CrHPRs in transcriptional level. Mean values ± SD are from three independent measurements. Means denoted by the same letter did not significantly differ at P<0.05.

Step 2: If CrHPRs function in photorespiration, they could be induced in such condition. Based on the assumption, we monitored their transcriptional profiles to determine the CrHPRs involved in photorespiration. As indicated in Fig. 1B, a 2-fold increasement of *CrHPR1* and *CrHPR4* was detected in their transcriptional levels when cells were transferred from CO_2_ to air conditions, suggesting their potential roles in photorespiration. By contrast, we did not observe significant difference of *CrHPR2* or *CrHPR5* transcripts between CO_2_ and air conditions, suggesting their fewer dominant roles in photorespiration than *CrHPR1* or *CrHPR4* (Fig. 1B).

Together, the NADH-dependent CrHPRs take parts in photorespiration as the major components, in which CrHPR1 and CrHPR4 may play more important roles than CrHPR2 and CrHPR5.

### Knockout of CrHPR1 impairs photorespiration

CrHPR1 was confirmed with high NADH-dependent activity, and may play major roles in photorespiration (Table 1, Fig. 1). To explore its physiological function, we set out to isolate the insertion strain for *CrHPR1*/*Cre06*.*g295450*.*t1*.*2* within the mutant library generated previously (Cheng et al., 2017). As shown in Fig. 2A, the paromomycin resistance cassette *AphVIII* was inserted into the seventh intron in *Crhpr1*, which was confirmed by the genome sequencing (data not shown). The knockout then was verified by qRT-PCR analysis using primers that span the cassette insertion site as described in methods. The mature transcript was found to be greatly disrupted in the mutant as indicated in Fig. 2B and C. These results confirmed the knockout of the *CrHPR1/Cre06*.*g295450*.*t1*.*2* in the insertion line *Crhpr1*, and used for the further study.

**Fig. 2.**
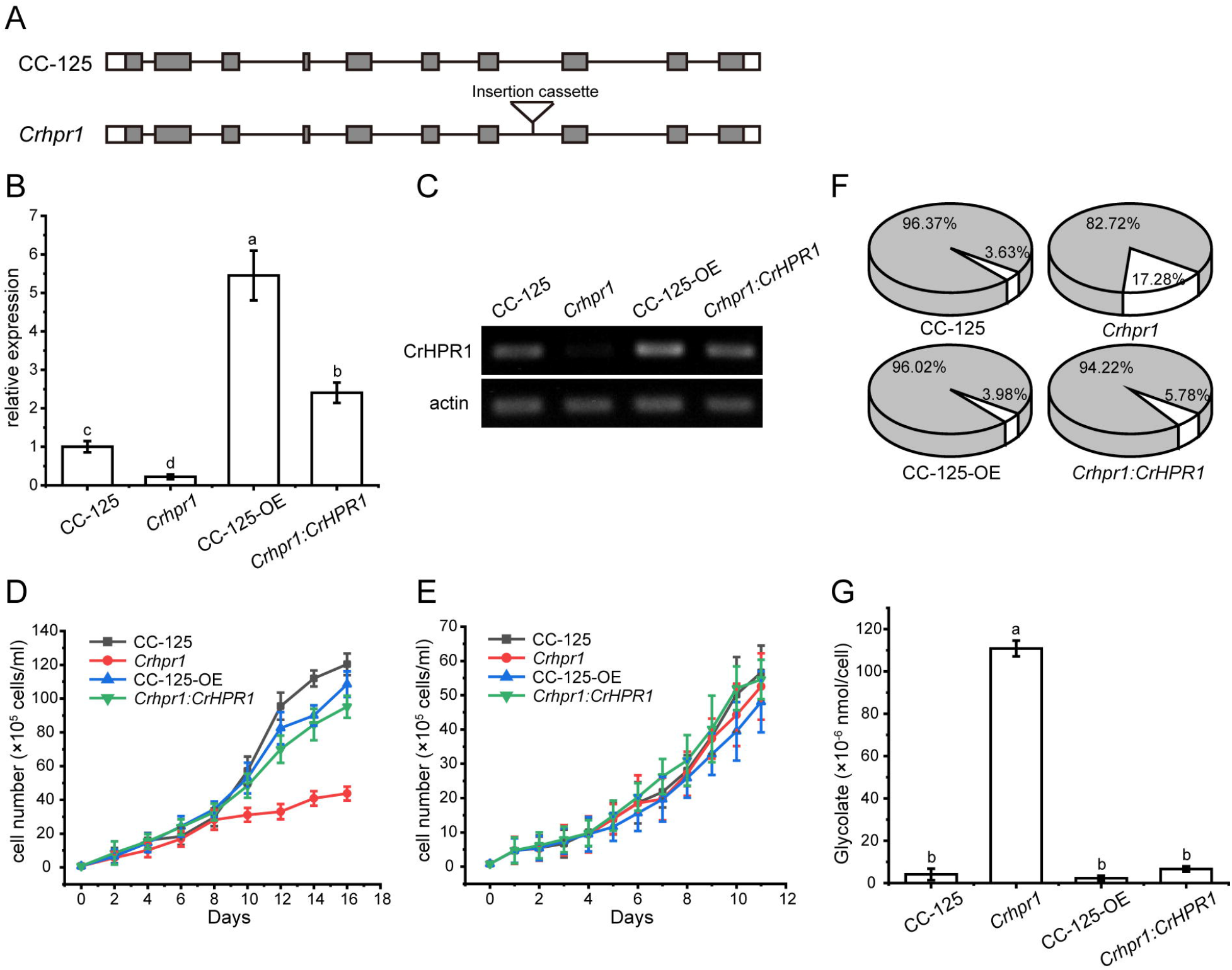
Knockout of *CrHPR1* impairs photorespiration. (A) Schematics of *CrHPR1* structures in WT/CC-125 and *Crhpr1*. (B) qRT-PCR analysis of *CrHPR1* transcripts in the strains. (C) RT-PCR of analysis of *CrHPR1* transcripts in the strains. (D-E) Growth curve of strains in air (D) and 3% CO_2_ condition (E). (F) The ratio of oxidation and carboxylation reaction of Rubisco in air. Carboxylation reaction: white sector, Oxidation reaction: gray sector. (G) Concentration of glycolate detected in the medium of each strain. Mean values ± SD presents data from three measurements. Means denoted by the same letter did not significantly differ at P<0.05.

As visible in Fig. 2E, *Crhpr1* presented no noticeable difference of growth to WT when they were cultured in tris-minimal medium under high CO_2_ conditions. However, the photorespiratory defects of *Crhpr1*, such as retarded growth and decreased ratio of Rubisco (oxidation/carboxylation), were observed when the cells were transferred to the air condition (Fig. 2D and F, Table S5). Interestingly, a stronger ability to process photosynthetic electron flow was detected in *Crhpr1*, manifesting as the elevation in the maximum photochemical quantum yield of PSII, maximum electron transfer efficiency, and reduction in the minimum saturating irradiance (Fig. S6). The phenotypic defects of *Crhpr1* may result from the leakage of photosynthetically fixed carbon, which is supported by the more glycolate excreted into the medium by *Crhpr1* as measured in Fig. 2G. Overexpression of *CrHPR1* in WT/CC-125 did not bring in visually phenotypic changes (Fig. 2D, E, F and G).

The phenotypes of *Crhpr1* detected above were restored to the WT/CC-125 level in the rescued strain *Crhpr1:CrHPR1* (Fig. 2B, C, D, F and G; Fig. S6), which provided evidence that the photorespiration is disrupted in *Crhpr1* and thus that the function of CrHPR1 is closely related to photorespiration.

### CrHPR2 knockdown strains show photorespiratory defects at *Crhpr1* background

Although CrHPR2 was detected with NADPH-dependent HPR activity, it was not induced in WT/CC-125 under photorespiratory condition as indicated by qRT-PCR analysis (Fig. 1A), and no difference in phenotypes of CrHPR2 knockdown strains and WT/CC-125 was observed (Table S6, Fig. S7). We speculated that CrHPR2 performs the compensatory functions of CrHPR1, and it may act as the major component in the extra-mitochondria hydroxypyruvate-reducing pathway in *Chlamydomonas*, which could catalyze the redundant hydroxypyruvate penetrated from mitochondria. To test this possibility, we compared the transcript level of *CrHPR2* in WT/CC-125 and *Crhpr1*. As shown in Fig. 3A, the expression of *CrHPR2* was greatly induced in air when *CrHPR1* was deleted, which verified the potential role of *CrHPR2* in photorespiration mentioned above. To explore its physiological function, we generated the *CrHPR2* knockdown strains at *Crhpr1* background, yielding serial *Crhpr1-a2* mutants. qRT-PCR analysis indicated that *CrHPR2* transcript in *Crhpr1-a2* strains was reduced to various extents of that in WT/CC-125 (Fig. 3B), which confirmed the disruption of *CrHPR2* expression. Thus, they were used for further analysis.

**Fig. 3.**
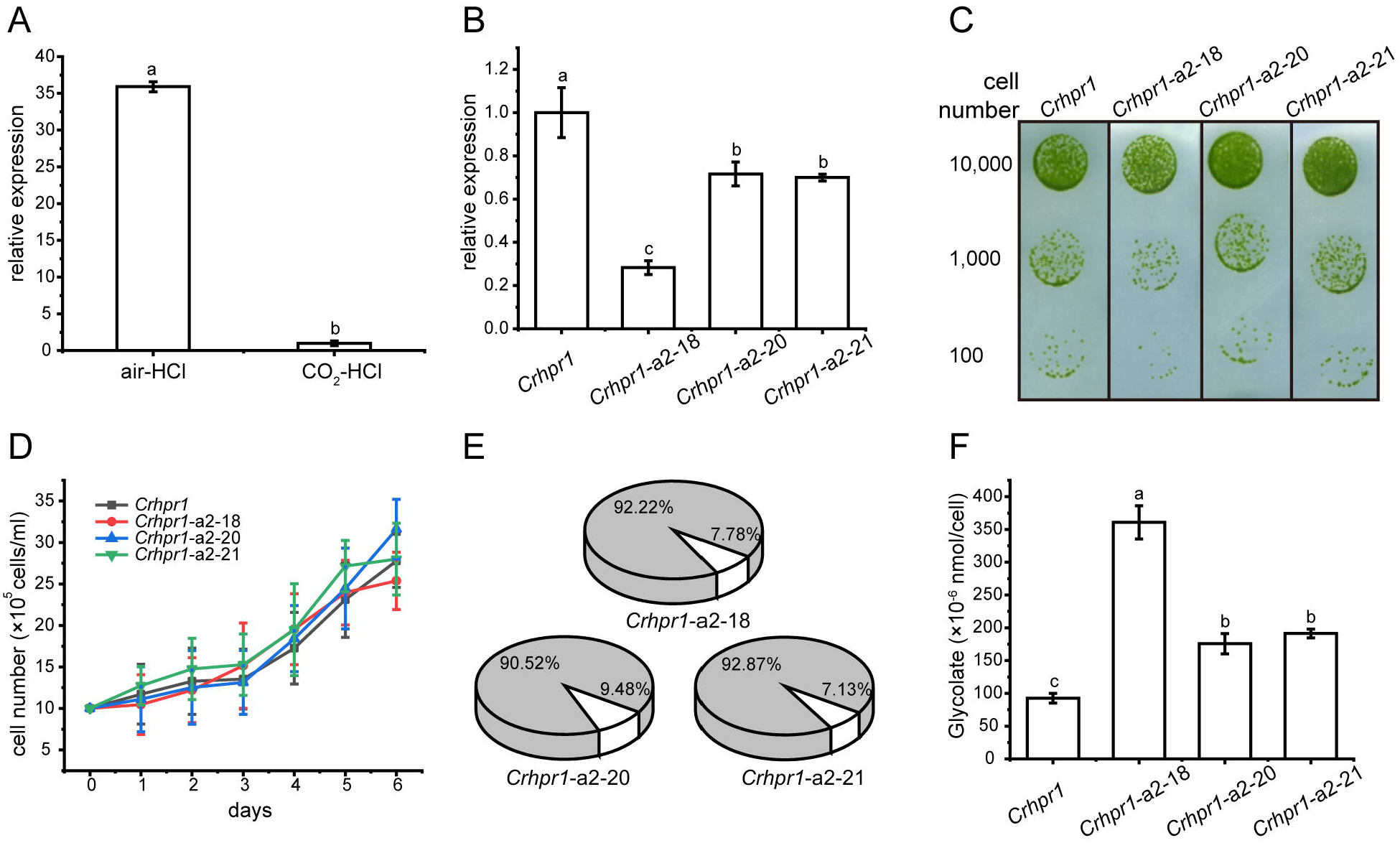
CrHPR2 knockdown strains show photorespiratory defects at *Crhpr1* background. A) Measurement of *CrHPR2* transcripts at both WT/CC-125 and *Crhpr1* background in air condition. (B) Measurement of *CrHPR2* transcripts in *Crhpr1* and *Crhpr1-a2* strains. (C) Spot tests showing growth of *Crhpr1* and *Crhpr1-a2* strains. (D) Growth curves of *Crhpr1* and *Crhpr1-a2* strains in air. (E) The ratio of oxidation and carboxylation reaction of Rubisco in *Crhpr1-a2* strains. Carboxylation reaction: white sector, Oxidation reaction: gray sector. (F) Concentration of glycolate detected in the medium of each strain. Mean values ± SD presents data from three measurements. Means denoted by the same letter did not significantly differ at P<0.05.

Although the expression of *CrHPR2* was greatly destroyed, no visible difference in growth was observed between *Crhpr1* and *Crhpr1-a2* strains when the cells were cultured in air, regardless of the solid or liquid medium (Fig. 3C and D). Nevertheless, the catalytic rate of Rubisco (carboxylation/oxidation) of *Crhpr1-a2* strains was much lower than that of *Crhpr1*, especially that the carboxylation rate was greatly affected in *Crhpr1-a2* (Fig. 3E, Table S7). These results suggest that the photosynthetic activity of *Crhpr1-a2* may have been affected by the increased glycolate excreted by the strains. We further measured the concentration of glycolate secreted to the medium by respective mutants, and the results indicated that more glycolate was indeed released by *Crhphr1-a2* strains than *Crhpr1* (Fig. 3F).

These evidences confirm that photorespiration is affected in the *CrHPR2* knockdown strains, and *CrHPR2* is indeed associated with photorespiration. This exacerbated excretion of glycolate by *Crhpr1-a2* may bring damage to photosynthesis in some extents, manifesting as the decreased quantum yield efficiency of PSII, electron transfer rate and minimum suturing irradiance (Fig. S8).

### CrHPR4 participates in photorespiration as a chloroplast-targeting glyoxylate reductase

*CrHPR4* was greatly induced in both WT/CC-125 and *Crhpr1* strains in air (Fig. 1A and 4A), demonstrating its participation in photorespiration. To uncover its detailed function in photorespiratory physiology, we generated the *CrHPR4* knockdown strains at WT/CC-125 background, yielding *WT/CC-125-a4*. Both photosynthetic activity and growth of *WT/CC-125-a4* strains were only slightly affected (Fig. S9), which is consistent with previous study (Burgess et al., 2015). However, the ratio of Rubisco (oxidation/carboxylation) was detected with a lower value in *WT/CC-125-a4* strains than that in WT/CC-125 (Table S8), suggesting the relation of CrHPR4 to photorespiration in some extents. To further explore the possibility, we generated the *CrHPR4* knockdown strains at *Crhpr1* background, yielding *Crhpr1-a4*. As determined by the qRT-PCR analysis, a reduced expression level of *CrHPR4* was detected in *Crhpr1-a4* strains (Fig. 4B) and they were used in the following studies.

**Fig. 4.**
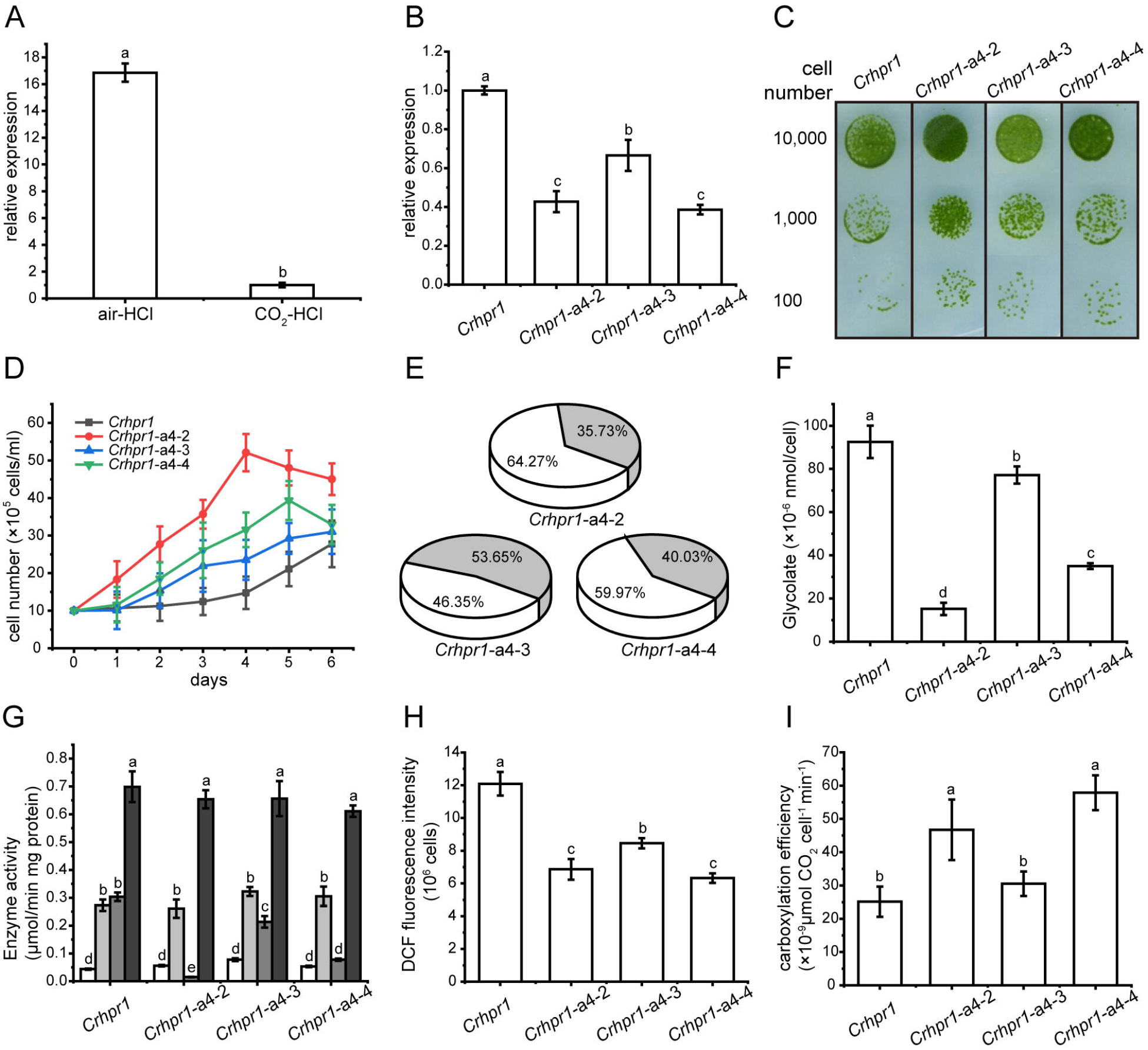
CrHPR4 participates in photorespiration as a chloroplast-targeting glyoxylate reductase. (A) Measurement of *CrHPR4* transcripts at both WT/CC-125 and *Crhpr1* background in air condition. (B) Measurement of *CrHPR4* transcripts in *Crhpr1* and *Crhpr1-a2* strains. (C) Spot tests showing growth of *Crhpr1* and *Crhpr1-a4* strains. (D) Growth curves of *Crhpr1* and *Crhpr1-a4* strains. (E) The ratio of oxidation and carboxylation reaction of Rubisco in *Crhpr1-a4* strains. Carboxylation reaction: white sector, Oxidation reaction: gray sector. (F) Concentration of glycolate detected in the medium of each strain. (G) Enzyme activity assay of *Crhpr1* and *Crhpr1-a4* strains. HPR-NADH: white bars, HPR-NADPH: light gray bars, GR-NADH: gray bars, GR-NADPH: dark gray bars. (H) Determination of ROS in *Crhpr1* and *Crhpr1-a4* strains treated with salicylhydroxamic acid (SHAM). (I) Determination of Carboxylation efficiency in *Crhpr1* and *Crhpr1-a4* strains treated with salicylhydroxamic acid (SHAM); Mean values ±SD from three measurements. Means denoted by the same letter did not significantly differ at P<0.05.

Apparently, the visually enhanced growth was observed for *Crhpr1-a4* strains compared to that of *Crhpr1* in both solid and liquid medium (Fig. 4C and D). The highly accumulated biomass of *Crhpr1-a4* possibly resulted from its efficient activity in both photosynthesis and CO_2_ fixation (Fig. 4E, S10; Table S9). Unexpectedly, less glycolate, intermediate in photorespiration, was excreted by *Crhpr1-a4* than both *Crhpr1* and *Crhpr1-a2* (Fig. 4F), and this prompted us to investigate whether CrHPR4 still perform functions as HPR. We then examined the enzymatic activity with cell extracts from respective strains, and determined the difference (Fig. 4G).

Intriguingly, the enzymatic activity of NADH-dependent glyoxylate reductase rather than HPR was significantly impaired/decreased in *Crhpr1-a4* strains compared to that in *Crhpr1* (Fig. 4G). These evidences established the relationship of CrHPR4 to photorespiration but it may mainly function as glyoxylate reductase not HPR, and its physiological effects seem to be different from those of CrHPR1 and CrHPR2.

Considering that CrHPR4 was targeted to the chloroplast (Fig. S5), it likely plays roles in the glycolate-quinone oxidoreductase system which could be inhibited by salicylhydroxamic acid (SHAM) (Goyal, 2002). To test this possibility, we added SHAM to the medium and found that the excretion of glycolate was indeed suppressed in *Crhpr1-a4 strains* (Fig. 4F). Moreover, the halted glycolate-quinone oxidoreductase reaction by SHAM resulted in less reactive oxygen species which is the byproduct of the oxidoreductase system (Fig. 4H), and higher carboxylation efficiency (Fig. 4I).

Together, these results provide the evidence that CrHPR4 may take parts in photorespiration by acting as the chloroplasdial glyoxylate reductase in the glycolate-quinone oxidoreductase system, which differentiates it from CrHPR1 and CrHPR2.

## DISCUSSION

### The ensemble of CrHPR proteins identified here is previously uncharacterized photorespiratory components with hydroxypyruvate reductase

In an effort to identify all the hydroxypyruvate reductase in Chlamydomonas genome, we initially used the BLAST function with the protein sequence of CrHPR1 as reference to search against the Chlamydomonas proteome. The analysis generated a list of 5 candidate novel CrHPRs (including CrHPR1 itself), in which the conserved domains of HPR, including NAD(P)-binding motif, NAD recognition sites, substrate-orienting and catalytic pair domains, were identified (Fig. S1 and S2). Further study of the five CrHPRs by enzymatic assay showed that all five are indeed detected with the activity of HPR (Table 1). Subcellular localization (Fig. S5) and phenotypic characterization of respective mutants (Fig. 2, 3 and 4) provide evidences that some of these candidate proteins in fact act in photorespiration. Thus, we have globally identified the CrHPRs and explore their functions that were previously unknown.

### CrHPRs are locating in multiple subcellular regions when function in photorespiration

Compared to the higher plants, photorespiration was assumed to pass through the mitochondria rather than peroxisome in Chlamydomonas (Nakamura et al., 2005), which is supported by the detected mitochondrial CrHPRs activity (Stabenau et al., 1974). Husic further demonstrated that it’s NADH-dependent CrHPRs that may function as the major components of HPRs in photorespiration (Husic and Tolbert, 1987), but not yet known at the sequence level. Here we identified two mitochondrion-targeting CrHPRs, CrHPR1 and CrHPR5, assayed with NADH-dependent activity for the first time according to our knowledge (Fig. S5, Table 1), and they are likely the proteins previously shown to be associated with the photorespiration (Husic and Tolbert, 1987). Especially CrHPR1, which was experimentally verified with high NADH-dependent enzymatic activity, may contribute most of the HPR activities measured by Husic (Husic and Tolbert, 1987). Therefore, it may not be coincidental that *Crhpr1* developed photorespiratory defects in air (Fig. 2 D, E, F and G). CrHPR5, active in the presence of NADPH, may have a function related to the glutathione peroxidase or peroxiredoxin antioxidant systems that need plenty of NADPH converted from NADH.

But Husic did not know that there are bypass pathways of photorespiration, a conclusion that is supported by both previous and present studies (Timm et al., 2008; Timm et al., 2011). In present study, we found that CrHPR2 and CrHPR4, which are proved to be related to photorespiration (Fig. 3 and 4), were targeted to cytosol and chloroplast (Fig. S5), respectively. They may function as such components of the bypass pathway of photorespiration, and it will be of interest to determine their detailed functions as discussed in the below sections.

### CrHPR1 acts as the major NADH-dependent HPR in Chlamydomonas, and functions differently from HPR1 in higher plants

CrHPR1 was assigned into the plant-specific subgroup in phylogenetics analysis (Fig. S4), and it has been well inherited through evolution implicated by the presence of homologs in higher plants (Fig. S1). However, the function of CrHPR1 in photorespiration may be different from its homologs in higher plants according to the present study.

Unlike the peroxisome-location of HPR1 in higher plants, CrHPR1 was targeted to the mitochondria (Fig. S5), which supports the assumption that photorespiration may pass through mitochondria rather than peroxisome in Chlamydomonas (Nakamura et al., 2005). Further investigation revealed that *Crhpr1* presented obvious photorespiratory defects and retarded growth in air (Fig. 2), which are greatly different to the no visually noticeable phenotypes of *hpr1* mutants in high plants (Timm et al., 2008; Cousins et al., 2011). This leads us to suggest that CrHPR1 acts as the major NADH-dependent HPR in Chlamydomonas, and the roles of CrHPR1 in photorespiration seems to be much more dominant than that of HPR1 in higher plants. Apparently, the cytosolic or chloroplast bypass pathway of photorespiration (Fig. 3 and 4), performed by CrHPR2 and CrHPR4, could not fully compensate the lost function of mitochondrial CrHPR1. Therefore, the novel phenotypes of *Crhpr1* described here expand the range of HPRs’ phenotypes associated with photorespiratory defects.

In the course of analysis, we also noticed that Chlamydomonas and high plants responses differently to the photorespiratory defects. In *Arabidopsis*, 2-phosphoglycolate is accumulated and metabolized by G6P shunt in *hpr1* mutant (Li et al., 2019), and hydroxypyruvate is converted into glycolate after decarboxylation and oxidation, then glycolate reenters the core photorespiration pathway (Missihoun and Kotchoni, 2018). However, the *Crhpr1* mutant excretes a large amount of glycolate to balance the internal environment (Fig. 2G), which resulted in the greatly reduced efficiency of carbon fixation. It’s likely that more complex photorespiration and adaption mechanisms have been adopted during evolution, but the underlying cause of the difference between Chlamydomonas and higher plants remains to be clarified.

### CrHPR2 participates in the cytosolic bypass of photorespiration

The general assumption that conversion of hydroxypyruvate to glycerate is exclusively performed by the mitochondria- or peroxisome-targeted HPR1 may not be comprehensive (Timm et al., 2008; Timm et al., 2011; Ye et al., 2014; Missihoun and Kotchoni, 2018), considering that HPR2 could participate in the cytosolic bypass of photorespiration in both Chlamydomonas and higher plants (Table 1, Fig. 3; Timm et al., 2008). Our determination of the functions of CrHPR2 in photorespiration is based on evidence from several aspects as follows.

In our initial analysis, we found that CrHPR2 was assigned into the bacterial subgroup, which suggests that its participation may not be limited to the light-related metabolism (Fig. S4). To further investigate its function, we generated the CrHPR2 knockdown strains at WT/CC-125 background, but no obvious photorespiratory defect was observed (Fig. S7). Only when CrHPR2 was knocked down in *Crhpr1*, the more severe photorespiratory defects of *Crhpr1-a2* could be detected than those of *Crhpr1* (Fig. 3). These evidences imply the participation of CrHPR2 in photorespiration, but it may play a compensatory role in some extent, which is supported by the result from qRT-PCR analysis (Fig. 3A). If so, these evidences mentioned above support the permeability of mitochondria matrix for hydroxypyruvate. Thus, our data provide indirect evidence that hydroxypyruvate could easily equilibrate with cytosol when no CrHPR1 is present within the mitochondria. Despite it remains to be examined the presence of suggested mitochondria channeling of photorespiratory intermediates HP (Keech et al., 2017), it is not unlikely that such equilibration mentioned above occurs in WT/CC-125.

Last but not least, it is of importance to build the interaction network of cytosolic CrHPR2, which would provide more details of CrHPR2 in both photorespiration and other metabolic pathways. What’s more, CrHPR3 was also localized to cytosol but did not show a similar function to that of CrHPR2. How CrHPR2 and CrHPR3 coordinate with each other in the cytosolic bypass of photorespiration remains to be explored.

### CrHPR4, targeted to the chloroplast, mainly plays roles in photorespiration as glyoxylate reductase within this compartment

Here, we report to our knowledge a previously uncharacterized enzyme that could reduce HP, glyoxylate and pyruvate (Table 1), and hence could directly or indirectly contribute to photorespiration in Chlamydomonas. Considering its multiple substrates and preference for the cofactor NADH, the identified CrHPR4 could support mitochondrial and cytosolic CrHPRs as well as chloroplastidial and cytosolic glyoxylate reductase (Ching et al., 2012; Brikis et al., 2017) for an optimal reduction of the respective intermediates (Fig. 1). Therefore, the observed effects of CrHPR4 on photorespiration clearly suggest its involvement in this process (Fig. 4).

By combining the data of enzymatic characteristics and the analysis of *Crhpr1-a4* strains, we proposed the detailed mechanism for *CrHPR4* participating in photorespiration as shown in Fig. 5: CrHPR4 was detected with the activity of glyoxylate reductase (Table 1), and it may act in the glycolate-quinone oxidoreductase system which is supported by the results presented in Fig. 5. When *CrHPR4* is knocked down in *Crhpr1*, excess glyoxylate is converted into CO_2_ which directly inhibits the oxidation reaction of Rubisco while promoting the carboxylation reaction (Fig. 4E, Table S10). Meanwhile, the production of glycolate mediated by CrHPR4 is greatly disrupted in *Crhpr1-a4* strains, resulting in the less excretion of glycolate into medium (Fig. 4F). Together, the knockdown of CrHPR4 results in the increased CO_2_ fixation and less loss of photosynthetic fixed carbon (Fig. 4H and I). As a result, *Crhpr1-a4* was observed with more robust growth compared to *Crhpr1* (Fig. 4C and D). Considering the great effects of CrHPR4 on the conversion of glyoxylate into glycolate, it may function as the dominant glyoxylate reductase in chloroplast. It’s undeniable that CrHPR4 have linked the photosynthesis and photorespiration closely via the glycolate metabolism, and the fine-tuning mechanism remains unidentified yet.

**Fig. 5.**
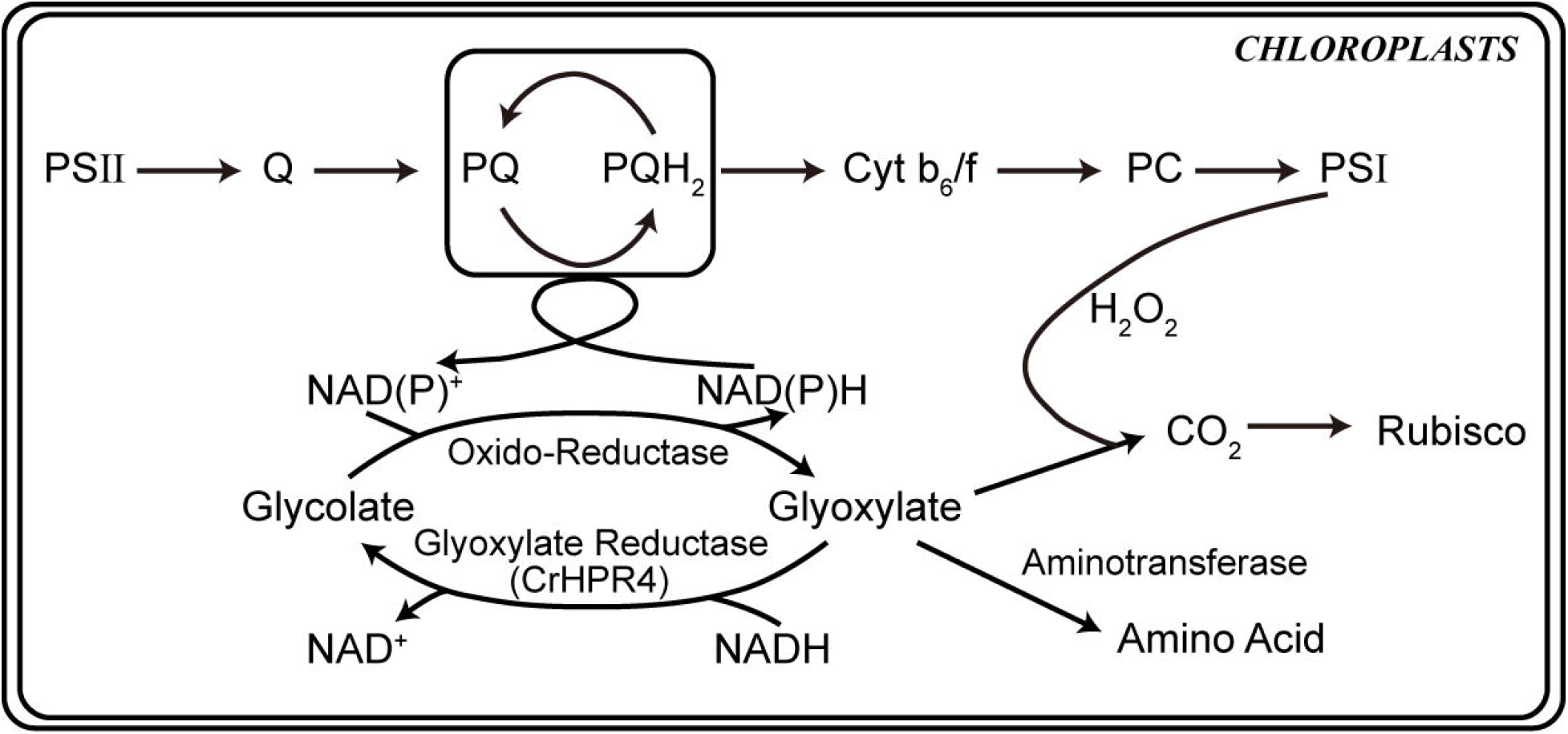
Schematics of the functional mechanism of CrHPR4 in photosynthesis and photorespiration.

Interestingly, CrHPR4 was detected with the activity of pyruvate reductase which is consistent with previous study (Burgess et al., 2015). As pyruvate reductase, CrHPR4, however, may mainly play an important role in anaerobic/fermentation metabolism (Burgess et al., 2015). It seems likely that CrHPR4 could participate in both photorespiration and anaerobic metabolism by acting as a bifunction enzyme, but how CrHPR4 performs roles in two pathways remains to be identified. Thus, the investigation of interaction network of CrHPR4 in chloroplast could provide more details. Moreover, CrHPR4 was also detected with the activity of HPR, but how HP is transported from mitochondrion to chloroplast is not identified yet (Reumann and Weber, 2006; Keech et al., 2017). Thus, it would be very informative to analyze and compare the “-omics” of *Crhpr1-a4* under photorespiratory and non-photorespiratory conditions as fermentation, to uncover its elusive functions and exact physiological role.

In conclusion, although the role for CrHPR4 will need further investigation, this study presents strong indications that the enzyme is closely associated with the photorespiratory process and can at least partially participate in the chloroplast glycolate metabolism. Moreover, CrHPR4 could display a possible link of photorespiration to photosynthesis and fermentation that remains to be identified. It’s particularly important to clarify the underlying mechanism of CrHPR4 functioning in multiple pathways, and it will benefit both the construction of plant metabolic network and the crop improvement by synthetic biology.

### Concluding remarks

Our results reveal that the CrHPRs are far more complex than previously recognized, and provide a greatly expanded knowledge base to understand their functions in photorespiration for future study. Considering the presence of multiple CrHPRs in Chlamydomonas genome and phenotypes of the mutants, it will be of great interest to uncover the detailed mechanism of how they coordinate with each other when performing roles in photorespiration.

CrHPRs could link photorespiration to photosynthesis and fermentation, and they may act as the central hub in the coordination of metabolism. Hence, the studies here will benefit the construction of plant metabolic network, and provide important clues for crop improvement by genetic engineering. Meanwhile, glycolate could be converted into methane (Günther er al., 2012), and it will be of interest to explore the potentiality of develop glycolate into bioenergy.

## MATERIALS AND METHODS

### Strains and culture conditions

All Chlamydomonas strains used in the work are listed in Table S1. WT strain used was CC-125 (Chlamydomonas Resource Center, https://www.chlamycollection.org). The mutant *hpr1* was generated as previously described (Cheng et al., 2017) and obtained from the Wu Han Jingyu Microalgae Science CO., LTD, China. AmiRNA mutants was generated as previously described (Hu et al., 2014).

Chlamydomonas cells were maintained in solid Tris-acetate-phosphate (TAP) plates, and cultured in liquid Tris-minimal medium (Gorman and Levine, 1965) at 25 °C under 80 μE m ^−2^ s ^−1^ continuous light. For high CO_2_ treatment, cells were grown in Tris-minimal medium with aeration of 3% CO_2_.

### qRT-PCR analysis

Total mRNAs used for qRT□PCR experiments were isolated from Chlamydomonas using the Eastep Super total RNA extraction kit (Promega), and cDNA was synthesized with the PrimeScript TMRT Master Mix kit (TaKaRa). qRT□PCR reactions were performed in triplicate using Mastercycler ep realplex (Eppendorf) with gene□specific primers and actin as the internal control (Winck et al., 2016). gene□specific PCR primer pairs used for the actin and CrHPRs are listed in Table S2. PCR primers were designed using Primer-BLAST of NCBI (https://www.ncbi.nlm.nih.gov/tools/primer-blast/index.cgi?LINK_LOC=BlastHome) and the amplifying program was as follows: pre-incubation at 95 °C for 30 s, 40 cycles of denaturation at 95 °C for 5 s, annealing at 62 °C for 5 s, and amplification at 72 °C for 25 s.

The change in fluorescence of SYBR Green I dye (SYBR Premix Ex Taq, TaKaRa) in every cycle was monitored by the realplex system software, and the cycle threshold (C_t_) above background for each reaction was calculated. The Ct value of actin was subtracted from that of the gene of interest to obtain aΔCt value. The Ct value of an arbitrary calibrator was subtracted from the ΔCt value to obtain a ΔΔCt value. The fold changes in expression level relative to the calibrator were calculated as 2^-ΔΔCt^.

### Gene cloning, heterologous expression of *Chlamydomonas* HPR and purification of recombinant proteins

Chlamydomonas cDNA samples were prepared as described above. HPR cDNAs were amplified using PrimeSTAR HS DNA Polymerase (Takara, Ohtsu, Japan) with gene□specific primers as listed in Table S3. The PCR product was cloned directly into pMD20-T vector (Takara, Beijing, China), which was then transformed into competent *E. coli* DH5α cells. Positive clones were verified by sequencing (BGI Genomics, Beijing, China).

For heterologous expression, the plasmids containing HPR2 were digested with *Nco*I and *Xho*I and the resulted fragment was linked to pETMALc-H treated with the same restricted enzymes (Kellyann et al., 1997), yielding pETMALc-H-HPR2. The plasmids containing CrHPR1, CrHPR3, CrHPR4 and CrHPR5 were digested with *Nde*I and *Xho*I, respectively, and then ligated into pET-30a digested with the same enzymes, yielding pET-30a-CrHPR1, pET-30a-CrHPR3, pET-30a-CrHPR4, pET-30a-CrHPR5. The constructed expression plasmids were transformed into expression host *E. coli* Rosetta (DE3) (CoWin Biosciences, Beijing, China), and the transformants were cultivated in the autoinduction medium ZYM2052 (Studier, 2005).

The recombinant HPRs was purified with HIS-Select nickel affinity gel filler (CoWin Biosciences, Beijing, China). Briefly, the supernatant of the broken cells collected and gently mixed with HIS-Select nickel affinity gel, and washed up by three cycles of binding buffer. The His-tagged HPRs was eluted with elution buffer.

### Generation of amiRNA, overexpression and CrHPR-CFP cell lines

For generation of amiRNA cell lines, pHK460 vector was given from Kaiyao Huang (Institute of hydrobiology, Chinese academy of sciences) (Nordhues et al., 2012). Target gene□specific oligonucleotide sequences were designed using the WMD3 software (http://wmd3.weigelworld.org). The resulting oligonucleotides that target CrHPR2 and CrHPR4 genes are listed in Table S4. Combined with the sequence of miRNA cre-MIR1157, these oligonucleotides were linked into pHK460 vector at the unique *Xho*I and *Eco*RI site. The plasmids were then isolated and subjected to transformation into Chlamydomonas cells.

For generation of overexpression cell lines, CrHPR1 were cloned from Chlamydomonas cDNA samples with gene-specific primers (*Crhpr1*-O-F/R) as in Table S3. The PCR product was connected into pMD20-T vector (Takara, Beijing, China) to form pMD20-CrHPR1, which was then transformed into competent *E. coli* DH5α cells. Positive clones were verified by sequencing (BGI Genomics, Beijing, China). The plasmid pMD20-CrHPR1 was digested with *Xho*I and *Eco*RI and the resulted fragment was linked to pHK460 treated with the same restricted enzymes. The mRNA from pHK460 is expressed from the hsp70/rbcs2 promoter and end at 3’UTR of /rbcs2. The transcript protein is fused with zeocin resistance selection maker, which will be cut off by the FMDV 2A self-cleaving sequence (Nordhues et al., 2012).

The constructed plasmids were transformed into Chlamydomonas cells using the electroporation method as described (Rasala et al., 2012). After transformation, cells were grown on Tris-acetate-phosphate agar supplemented with 15 μg/mL zeocin (Sigma-Aldrich). Colonies derived from single cells were picked for DNA extraction, after digested with RNase, Colonies were confirmed by PCR with primers (*Crhpr1*-O-F/R) to amplify the stripe which length is same as cDNA of CrHPR1.

For generation of CrHPR-CFP cell lines, CrHPRs were cloned from Chlamydomonas cDNA samples with gene-specific primers (CrHPRs-NT-F/R for testing N-terminal target signal and fuse the CFP to C-terminal of CrHPRs; CrHPRs-CT-F/R for testing C-terminal target signal and fuse the CFP to N-terminal of CrHPRs) as in Table S3. The PCR products were connected into pMD20-T vector (Takara, Beijing, China) to form pMD20-CrHPRs-NT or pMD20-CrHPRs-CT, and transformed into competent *E. coli* DH5α cells. Positive clones were verified by sequencing (BGI Genomics, Beijing, China). The plasmid pMD20-CrHPRs-NT was digested with *Xho*I and *Bam*HI, and the plasmid pMD20-CrHPRs-CT was digested with *Bam*HI and *Eco*RI, then the resulted fragment was linked to pHK460 treated with the same restricted enzymes.

The plasmids were then isolated and subjected to transformation into Chlamydomonas cells. Transformants were selected from TAP plates supplemented with 15 μg/mL zeocin (Sigma-Aldrich).

### Fluorescence microscopy

Representative cells were collected from TAP plates supplemented with zeocin. Images were captured on Image-Pro Express 6.0 (Media Cybernetics, Rockville, MD, USA) using Olympus BX51 (Center Valley, PA, USA) with Retiga-2000R camera (QImaging, Tucson, AZ, USA). Filters used in this research are CFP (excitation 436/10 nm, emission 470/30 nm) and Chloro (excitation 500/23 nm, emission 535/30 nm). The fluorescence images were false-colored using Adobe Photoshop CS3.

### Enzymatic activity assays

Cells were collected and broken in 1 mL extraction buffer (10 mM Tris-HCl, 1 mM EDTA, 2 mM MgCl_2_, 1mM β-mercaptoethanol, pH 7.5), and the supernatant was collected for the determination of enzymatic activity after centrifugation at 4 □. With 0.2 mM NADPH, 0.5 mM hydroxypyruvate or 1 mM glyoxylate, and 50 μL purified enzyme in 200mM sodium phosphate buffer (pH 6.5), The hydroxypyruvate and glyoxylate reductase activity was determined according to published procedures (Husic and Tolbert, 1987) by measuring the absorbance of NAD(P)H at 340 nm using Microplate Photometer (Thermo-FC) at 25 °C. The protein concentration was measured by Brandford method.

### Chlorophyll fluorescence measurement

Cells were immobilized, aiming to acquire accurate data, as described by Luz with modification (Luz et al., 2015). Briefly, cells were cultured in Tris-minimal medium, and collected when reached to the late logarithmic phase. Then the cells were washed with 0.85% (w/v) NaCl, and concentrated by 12 times. 2 mL of the concentrated cells were mixed with 8 mL of 2% sodium alginate solution. The 4 cm^2^ squares of monofilament nylon with 24 threads per inch was sterilized, and immersed into the above mixture. The monofilament nylon square was quickly transferred into 2% CaCl_2_ solution to solidify sodium alginate. The monofilament nylon squares were rinsed with 0.85% NaCl to remove excess solution. The above steps were repeated once again to get two layers of cell immobilized sodium alginate onto the squares.

The immobilized cells were clipped with DLC-8 dark leaf clip and measured by MINI-PAM-II (Walz, Germany) with the saturation pulse method by following the manufacturer’s instruction (Klughammer and Schreiber, 1994). Briefly, with the saturation pulse technique, the maximum quantum efficiency of PS□ (Fv/Fm) was detected after 30 min dark acclimated. The light curves were generated by using increasing actinic irradiance sequence ranging from 0 to 500 μmol photons m^-2^s^-1^.

### Measurement of photosynthetic and respiration rates

Cells at late logarithmic phase (3∼5×10^6^ cells/mL) were collected and concentrated by 5 times. The oxygen exchange was measured with a Chlorolab-2 oxygen electrode (Hansatech, Norfolk, UK) at 30 °C under both dark and light conditions by following the manufacturer’s instructions. Light intensity was determined using a quantum photometer (Hansatech).

The electron flow of Rubisco was calculated by combining the data of oxygen exchange and chlorophyll fluorescence (Valentini et al., 1995). Formula: Jc =1/3[ETRII + 8(A +R_D_)], Jo = 2/3[ETRII -4(A+ R_D_)], Jc: the electron flux of Rubisco carboxylation; Jo: the electron flux of Rubisco oxygenation; A: net photosynthetic rate; R_D_: day respiration.

### Quantitative measurement of ROS

The ROS of cells were measured by Reactive Oxygen Species Assay Kit (Beyotime Institute of Biotechnology, Shanghai, China) following the manufacturer’s instructions. Briefly, Cells at late logarithmic stage were collected, and stained with 100 μM 2’,7’-dichlorofluorescein diacetate (H_2_DCF-DA) for 45 min at room temperature as described previously (Affenzeller et al., 2009). Then cells were washed 3 times with Tris-minimal medium to remove the unbound probes and resuspend in 1mL Tris-minimal medium. The cells mixture was measured by fluorescence spectrophotometer (Enspire, PerkinElmer LLC, US) by using 485 nm excitation and 530 nm emission.

### Glycolate Determination

Cells were collected at late logarithmic phase and added to new Tris-minimal medium to make the final concentration reach at 2×10^6^ cells/mL. After 48 hours, glycolate was detected from the supernatant according to Kenji’s method (Takahashi, 1972). Samples containing 0.1 to 10 μg glycolate were used to draw the standard curve. 50 μL samples were added to the test tube with 1mL 0.01% 2,7-dihydroxynaphthalene in concentrated sulfuric acid. After 20min of 100□ water bath, the mixture was measured by Microplate Photometer (Thermo-FC) at 540 nm.

### Bioinformatic analysis

To identify the Chlamydomonas HPR proteins, CrHPR1 protein sequence was searched against the *C. reinhardtii* predicted protein database in Phytozome 12 using the BLASTP function, searched against the HMMER *C. reinhardtii* reference proteome using the HMMER website service (https://www.ebi.ac.uk/Tools/hmmer/search/phmmer) with the default parameters, or searched against AlgaePath http://algaepath.itps.ncku.edu.tw/algae_path/home.html). Protein alignments were performed using ClustalW (Larkin et al., 2007) and viewed using the GeneDoc software (Nicholas et al., 1997). The maximum likelihood phylogenetic tree was produced using the MEGA 7 program (Kumar et al., 2015).

## ACKNOWLEDGMENTS

We are grateful to Dr. K.Y. Huang from the Institute of Hydrobiology, Chinese Academy of Sciences for his generous gift of plasmid and technical guidance of electroporation method.

This work was supported by grant from Tianjin Synthetic Biotechnology Innovation Capacity Improvement Project (TSBICIP-CXRC-027) to L.Z.

The authors declare no competing financial interests.

## AUTHOR CONTRIBUTIONS

M.L.S., L.Z. and Y.W. conceived the project; M.L.S. performed the experiments; M.L.S., L.Z. and Y.W. analyzed the data, wrote, and revised the manuscript; All the authors have read and approved the manuscript prior to submission.

## Supplemental Data

The following materials are available in the online version of this article.

**Table S1**. All Chlamydomonas strains used in the work.

**Table S2**. Oligonucleotides used in qRT□PCR.

**Table S3**. Oligonucleotides used in gene cloning.

**Table S4**. Oligonucleotides of amiRNA that target *CrHPRs*.

**Table S5**. Oxidation/carboxylation rate of Rubisco of WT/CC-125, *Crhpr1* and the rescued strains in air.

**Table S6**. Oxidation/carboxylation of Rubisco of CC-125-a2 strains in air.

**Table S7**. Oxidation/carboxylation of Rubisco of *Crhpr1-a2* strains in air.

**Table S8**. Oxidation/carboxylation of Rubisco of CC-125-a4 strains in air.

**Table S9**. Oxidation/carboxylation of Rubisco of *Crhpr1-a4* strains in air.

**Fig. S1**. Sequence alignment of hydroxypyruvate reductase homologues from select species.

**Fig. S2**. Sequence alignment of *Chlamydomonas* hydroxypyruvate reductase.

**Fig. S3**. Expression and purification of recombinant CrHPRs assayed by SDS-gel.

**Fig. S4**. Phylogenetic tree of HPR proteins inferred by bacterial and eukaryotic sources.

**Fig. S5**. Subcellular localization of the CFP reporter fused with N- or C-terminal peptides from CrHPRs

**Fig. S6**. Measurement of photosynthetic activity of *Crhpr1* and the rescued strains by chlorophyll fluorescence.

**Fig. S7**. Phenotypic analysis of the *CrHPR2* knockdown strains at CC-125 background.

**Fig. S8**. Measurement of photosynthetic activity of *Crhpr1-a2* strains by chlorophyll fluorescence.

**Fig. S9**. Phenotypic analysis of the *CrHPR4* knockdown strains at CC-125 background. **Fig. S10**. Measurement of photosynthetic activity of *Crhpr1-a4* strains by chlorophyll fluorescence.

